# Reduced palatability, fast flight, and tails: Decoding the defence arsenal of Eudaminae skipper butterflies in a Neotropical locality

**DOI:** 10.1101/2023.12.20.572531

**Authors:** Daniel Linke, Jacqueline Hernandez Mejia, Valery N. P. Eche Navarro, Letty Salinas Sánchez, Pedro de Gusmão Ribeiro, Marianne Elias, Pável Matos-Maraví

## Abstract

1. Prey often relies on multiple defences to avoid predators, such as difficulty of capture, attack deflection, or unpleasant taste. Eudaminae skipper butterflies rely on behavioural (fast flight) and mechanical defences (hindwing tails), but other defences like unpalatability (consumption deterrence), and the interaction among defences, have not been assessed.
2. We test the palatability of 12 abundant Eudaminae species from a Neotropical locality, using training and feeding experiments with domestic chicks. Further, we approximate the difficulty of capture explained by flight speed and quantified by wing loading. To test how multiple defences are associated and explained by body size and habitat preference, we perform phylogenetic Bayesian regression analyses.
3. We found a broad range of palatability in Eudaminae, within and among species. Palatability was negatively correlated with wing loading, indicating that butterflies with higher flight speed tend to have lower palatability.
4. The presence or absence of tails did not explain the level of butterfly palatability, showing that attack deflection and consumption deterrence are not mutually exclusive defences. Habitat preference (open or forested environments) did not explain the level of palatability in Eudaminae, but it seems that butterflies with high wing loading (fast flight) tended to occupy semi-closed (e.g., hilltops) or closed habitats (e.g., secondary forest).
5. Finally, the level of unpalatability in Eudaminae is size dependent, larger butterflies are less palatable, perhaps due to their more likely detectability/preference by predators. Altogether, our findings shed light on the contexts favouring the prevalence of single vs. multiple defensive strategies in prey.

## Introduction

Many prey species simultaneously exhibit multiple anti-predator defences, which are often displayed at different stages of the predator-prey-interaction (i.e., prey encounter, detection, identification, attack, capture) (Ruxton et al., 2018). For example, to reduce detection by predators, visual defences such as crypsis are more successful when expressed in prey with smaller body size (Kang et al., 2017; Pembury Smith & Ruxton, 2021). Upon detection and approach by predators, prey may further increase their survival chance via different behavioural defences (e.g., escape ability and burst speed), mechanical defences (e.g., morphological structures that draw attacks towards non-vital body parts), or typically a synergy of both (Ruxton et al., 2018). At the post-contact predation sequence, chemical defences producing unpalatability might successfully deter prey consumption and are frequently associated with visual cues that act as warning signals (i.e., aposematism). In nature, all anti-predator strategies are imperfect, as predators evolve counteracting mechanisms (e.g., tolerance to chemical defences; Fink & Brower, 1981) and all ecosystems contain multiple types of potential predators (Guariento et al., 2022). Thus, a major question in evolutionary ecology is how often anti-predator defences transition within lineages over evolutionary time (e.g., Loeffler-Henry et al., 2023), and which contexts favour single-defence specialization versus multiple anti-predator defences (e.g., McClure et al., 2019).

The co-expression of multiple defences is shaped by trade-offs in resources and functions as well as interactions between traits where some defences may mutually benefit or constrain each other (Kikuchi et al., 2023; Ruxton et al., 2018). In Lepidoptera, one of the best-studied groups in terms of multi-defence evolution, such trait interactions result in certain prevalent defence combinations. For example, visual and behavioural defences are often co-exhibited, such as the cryptic ventral side of the Peacock butterfly (Nymphalidae: *Aglais io*) resembling dead leaf matter, along with their large eyespots on the dorsal side that are briskly displayed and accompanied with a hissing sound, thereby scaring predators off and facilitating escaping (Vallin et al., 2006, Olofsson et al., 2012). Likewise, Ithomiini butterflies have evolved partly transparent wings, which makes them concealed from predators (Arias et al., 2019), while being at the same time unpalatable and aposematic (McClure et al., 2019; Pinna et al., 2021). While all these defences increase the fitness of their bearer, our understanding is limited on whether different types of defences are associated with different habitats, or whether they have evolved due to different selective pressures along the predation sequence (Exnerová et al., 2023).

Studies of multiple anti-predator defence evolution in tropical butterflies have mostly focused on chemically defended and visually aposematic species. Our work focuses on a hyper diverse lineage of Neotropical butterflies that largely rely on behavioural and mechanical defences, though visual and chemical defences may still be part of their defence strategies. We study co-existing Eudaminae skipper butterflies (Lepidoptera: Hesperiidae, ca. 560 described species), while focusing on the most abundant species found at the foothills of the northeastern Peruvian Andes. Skippers exhibit robust thoracic muscles that support their manoeuvrable flights and elusive behaviours (Betts & Wootton, 1988; Sourakov, 2009, 2011). Thus, their fast flight (behavioural defence) likely increases their chances of survival at the pre-contact predator-prey-interaction sequence. Skippers might indeed be associated with high flight speeds compared to other butterflies, and this can be predicted by their high wing loading, i.e., body mass relative to wing area (Le Roy et al., 2019). Furthermore, Eudaminae skippers possess hindwing tails, potentially deflecting attacks towards non-vital body parts (mechanical defence; as in, e.g., swallowtail butterflies, Chotard et al., 2022). However, it remains unclear whether the repeated evolution of hindwing tails in different Eudaminae species (Li et al., 2019) was driven by predatorial selection or other factors. Finally, despite lacking typical warning colourations, certain Eudaminae phenotypes, such as the green-blueish iridescence on dorsal wings, may represent honest cues signalling unprofitability that have evolved multiple times in unrelated lineages (Janzen et al., 2009). However, unprofitability in the form of unpalatability (chemical defence) has not been thoroughly investigated in Eudaminae, as these have generally been assumed to be palatable due to their fast flight and escape capabilities (Pinheiro, 1996).

We hypothesize that the abundance of butterflies with various defences (behavioural, mechanical, and potentially chemical) can be explained by two factors: 1) by different pressures along the predator-prey-interaction sequence or 2) by habitat association wherein multiple types of predators may co-exist, potentially selecting for multiple types of prey defences. In this study we quantify defences of Eudaminae species, namely **1) behavioural defences**, as individual wing loading, which is positively linked to increased flight performance and speed, and can be regarded as a robust behavioural proxy in butterflies (Srygley & Chai, 1990; Berwaerts et al., 2002; Le Roy et al., 2019), including skippers (Dudley, 1990; Dudley & Srygley, 1994) **2) mechanical defences**, as species-level hindwing tails presence or absence; and, if found, **3) chemical defences**, as any individual variation of palatability.

We assess the following predictions in a correlational phylogenetic framework:

I. If Eudaminae butterflies do not possess chemical defences, we predict a positive correlation between the presence of hindwing tails (a mechanical defence) and wing loading indicative of fast flight and escape ability (a behavioural defence). This alignment is commonly observed in Lepidoptera (Kikuchi et al. 2023).
II. If certain Eudaminae species possess chemical defences evidenced by reduced palatability, we expect them to show low wing loading compared to Eudaminae that do not possess chemical defences, as chemically defended butterflies are typically associated with slow movement and low wing loading (Marden & Chai, 1991; Pinheiro, 1996; Srygley, 1994, 2004; but see Pinheiro et al., 2016).
III. Further, following a functionally non-redundant scenario outlined by Kikuchi et al. (2023), unpalatability (consumption deterrence) and hindwing tails (attack deflection) are expected to be negatively correlated because both defences operate at the post-contact predator-prey-interaction sequence.
IV. If such expectations are not met for certain Eudaminae butterflies with low palatability, we predict that the expression of chemical defences might be more beneficial, and therefore more commonly found, in large individuals and species, due to size-dependent detectability or preference along the predation sequence; e.g., size-dependent behavioural (Dewitt et al., 1999) and visual defence evolution (Pembury Smith & Ruxton, 2021).
V. Alternatively, species expressing chemical defences might be associated with different types of habitats (open/disturbed or closed/semi-open).

## Materials and methods

### Butterfly collection

The study location was at Wayrasacha, San Martin region, Peru (6.461 S, 76.334 W), where secondary growth forest is interspersed with agricultural areas, near the “Área de Conservación Regional Cordillera Escalera”. We collected the most abundant Eudaminae species at the study site: *Cogia crameri* (Oileidini: Typhedanina), *Drephalys helixus* (Entheini), *Cecropterus dorantes*, *C. doryssus*, *C. longipennis*, *C. zarex*, *Spicauda simplicius*, *Urbanus* spp., *Telegonus fulgerator*, *Epargyreus* spp. and *Chioides catillus* (Eudamini: Eudamina). As a negative reference (unpalatable butterfly), we used the abundant *Heliconius melpomene* (Nymphalidae: Heliconiinae).

Butterflies were captured with hand nets. Most butterflies were identified to the species level in the field, and all specimens were molecularly identified a posteriori in the lab (see below). Butterflies were stored alive in glassine envelopes with wet paper to avoid desiccation until the palatability experiments. The surrounding habitat (∼5 m) of every caught individual was determined as “open/disturbed” (open ground, grassland, shrubs) or “closed/semi-open” (forest edge, secondary or primary forest). As a conservative estimate of habitat association, given the high mobility of skippers and to account for stochastic events, we assigned to each species either habitat category based on the majority of habitat occurrences. We also revised the literature to assess any significant departure in habitat associations of our study species in other Neotropical localities (e.g., Janzen et al., 2011; Paluch et al., 2016; Zhang et al., 2023).

### Butterfly palatability

Butterflies were kept alive in humid conditions to avoid desiccation for a maximum of 24 hours and killed no more than 10 minutes before the start of the experiment to avoid any tissue and chemical degradation. The wings were removed and stored for subsequent morphometric analyses. Three legs per specimen were stored in 90% ethanol for subsequent molecular species identification.

We then prepared testing pellets consisting of crushed butterfly body and chick starter (Corimix, Corina Alimentos, Lima, Peru), as in Chouteau et al. (2019) and McClure et al. (2019). As certain skipper individuals were difficult to identify in the field, especially those with damaged wings, the palatability tests were conducted for each individual separately, rather than pooling individuals as in McClure et al., (2019). The butterfly body was weighted using a TGD 50-3C balance (precision 0.001 g, Kern & Sohn GmbH, Balingen, Germany). This measurement will later be used as body mass (representing the total weight of the butterfly without wings and three legs). After weighting and adding the same amount of chick starter, we added two drops of food colourant (Fratello, Importaciones Goicochea SAC, Lima, Peru) and crushed all together in clean stone mortars. The dye colour was alternated between orange and green for every other butterfly specimen per species, to rule out any colour preference or any association of a particular colour with (un)palatable meals by the chick. As palatable reference, we used mealworms (*Tenebrio molitor*) and as unpalatable reference *Heliconius melpomene*. As experimental control pellets (for each experiment), we used only chick starter in amounts similar as the testing pellets (either butterfly or mealworm). All the resulting pellets weighted ∼0.2–0.8 g.

For the most abundant species in the study area, *Spicauda simplicius*, we assessed whether removing the wings might affect the butterfly palatability test. In such cases, we photographed the left and right sides of the butterfly before pellet preparation. Results from this experiment can be found in Supplement 4.

To test the palatability of butterflies, we used domestic chicks (*Gallus gallus domesticus*) as surrogate avian predators (e.g., Chouteau et al., 2019; McClure et al., 2019). The palatability experiments were performed in experimental plywood cages of 60 x 59 x 59 cm (see Supplement 1) and consisted of three phases: (**1**) pre-training in group, (**2**) individual pre-training, and (**3**) palatability measure. Each chick was used only once in the palatability measure phase (**3**). All chicks had constant access to water and were accompanied with a “buddy” chick placed in a separate, contiguous cage to reduce stress.

1. Pre-training in group: To facilitate the initial learning and to speed up the screening process of trainable chicks, we used three to five, nine-day-old chicks in this stage. Chicks were starved for 60 minutes and were then placed in the experimental cage along with two experimental control pellets (∼0.5 g), green and orange. These were offered for a maximum of ten minutes. We considered the chicks successfully pre-trained once they consumed the two pellets in five consecutive rounds.
2. Individual training: The pre-trained chicks were starved for 60 minutes and were then individually offered two experimental control pellets, green and orange. We considered the individual chick fully trained when it completely consumed both pellets within a 10-minute frame.
3. Palatability measure: The fully trained chicks were 10 to 14 days old. After 60 minutes of food deprivation, the chick was left in the cage for ten minutes to destress, and afterwards, an experimental control and a testing pellet –butterfly (skipper or *H. melpomene*) or mealworm– with different colours were offered on separate metal coasters for a maximum of 10 minutes; the colours were alternated for each butterfly species. Chick behaviour, including picking pellets, water drinking, and beak cleaning, was noted as timestamps. We considered the palatability measure valid when the chick consumed more than 50% of the experimental control pellet and picked or approached extremely close the testing pellet at least once.

All experiments were filmed, and exemplary videos for each experimental phase can be found in the Supplementary Material 2.

### Molecular species Identification

DNA was extracted from three butterfly legs using QIAGEN kits and following the manufacturer’s instructions. We used Eudaminae-specific primers to amplify the mitochondrial COI barcoding region (see Supplement 3). DNA was sequenced by the company Macrogen (Amsterdam, Netherlands) and the chromatograms and alignments were processed using the software Geneious Prime v. 2023.0.1 (www.geneious.com). Specimens were identified using the COI barcodes and the BOLD database (www.boldsystems.org), and the species identities were further corroborated using the stored wings.

Because we rely on a single locus, we constrained the relationships and divergence times among genera following published phylogenomic backbones containing all the study Eudaminae genera (Li et al., 2019; Kawahara et al., 2023). We inferred a time-calibrated phylogenetic tree using MrBayes v. 3.2.6 (Ronquist et al., 2012) and IQ-TREE v. 2.1.3 (Minh et al., 2020) via the least square dating method (To et al., 2016) (Supplementary Material 3).

### Morphometric analysis

We scanned the dorsal side of the detached wings using a Color LaserJet Pro MFP M277dw printer (HP, Palo Alto, USA). We measured conventional wing morphometrics using the forewings of every butterfly individual; hindwings were typically damaged, especially those species with hindwing tails, thus, these were conservatively excluded from our morphometrics analysis. The measurements were taken using GIMP version 2.10.24 (GIMP Development Team, 2019) from the most intact forewing. Wing loading was calculated as the ratio of body weight relative to the area of both forewings.

### Statistical analyses

All statistical analyses were done in R v. 4.3.0 (R Core Team, 2022) and the graphics were generated using the package *ggplot2* v. 3.4.2 (Wickham, 2016).

To determine the level of butterfly palatability, we calculated a consumption ratio (*Cr*) for every butterfly specimen:

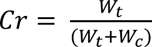

where, W_t_ is the proportion of consumed testing pellet (consumed weight relative to the initial pellet weight) and W_c_ is the proportion of consumed experimental control pellet. A *Cr* value close to 0.5 indicates high butterfly palatability while *Cr* close to 0 indicates a strong butterfly unpalatability. *Cr* values above 0.5, indicate the chick ate more of the butterfly testing pellet than the experimental control pellet.

To our surprise, we found a large variation in palatability levels between and within species. A Shapiro-Wilk normality test revealed non-normal distribution for *Cr*. We thus performed a pairwise Wilcoxon signed-rank test to compare the distribution of *Cr* between all distinct species with Bonferroni correction applied for multiple comparisons. Note that this species-level categorization does not influence the individual-based tests of our main predictions below. To test the phylogenetic signal of *Cr*, Pagel’s lambda and Bloomberg’s K were calculated for the species level averages and medians using the ‘phylosig’ function of *phytools* v. 1.5-1 (Revell, 2012). As we could not conclusively reject the null hypothesis of no phylogenetic signal, phylogenetic relatedness were included in the following Bayesian regression models.

We evaluated our predictions on the prevalence of certain defence combinations and the association of multi-defended prey with body size and/or habitat, using phylogenetic Bayesian models. We removed the references *Heliconius melpomene* and *Tenebrio molitor* from our dataset and we fitted five different Bayesian MCMC (Markov chain Monte Carlo) regression models, with 4 chains and 4,000 iterations each, using the R package *brms* v. 2.19.2 (Bürkner, 2017). To account for phylogenetic relatedness among species, the species identity of every individual and a variance-co-variance matrix calculated from our time-calibrated phylogeny using the R package *ape* v. 5.7-1 (Paradis & Schliep, 2019) were fed to the regression model as random factors. A description of all tested models (response and predictors) including the full model syntax can be found in Table 1.

**Table 1:** Summary of all fitted Bayesian regression models, including the tested predictions, used responses and predictors variables, and the full model syntax. Model (IV) was split into (IVa) and (IVb) as collinearity between body weight and forewing length might impact the stability and interpretability of regression models.

| Tested prediction | Response | Predictors | Model syntax |
| --- | --- | --- | --- |
| (I) Positive correlation between flight speed | Wing loading (WL; proxy for flight) | Hindwing tails (presence/absence; mechanical defence) | WL ~ tail + (1 \| gr(species_genetic, |
| and the presence of hindwing tails. | prowess/speed; behavioural defence) |  | cov = GenDist)) + (1 \| species) |
| **(II)** Specimens with chemical defences are associated with slow movements. | Consumption ratio (*Cr*; palatability measure) | Wing loading | $Cr \sim WL + (1 \vert \text{gr}(\text{species\_genetic}, \text{cov} = \text{GenDist})) + (1 \vert \text{species})$ |
| **(III)** Negative correlation between attack deflection (hindwing tails) and consumption deterrence (decreased palatability). | Consumption ratio | Hindwing tails (tail), interacting with forewing length (FW), as the functional effect of tails may depend on the size of forewings (Linke et. al *in prep*) | $Cr \sim \text{tail} * FW + (1 \vert \text{gr}(\text{species\_genetic}, \text{cov} = \text{GenDist})) + (1 \vert \text{species})$ |
| **(IVa)** Heavier specimens tend to be less palatable | Consumption ratio | Butterfly body weight | $Cr \sim \text{weight} + (1 \vert \text{gr}(\text{species\_genetic}, \text{cov} = \text{GenDist})) + (1 \vert \text{species})$ |
| **(IVb)** Larger specimens tend to be less palatable | Consumption ratio | Forewing length (FW) | $Cr \sim FW + (1 \vert \text{gr}(\text{species\_genetic}, \text{cov} = \text{GenDist})) + (1 \vert \text{species})$ |
| **(V)** Palatable butterflies are associated with open environments and less palatable butterflies are associated with forested environments | Consumption ratio | Average habitat for the species (habitat_sp), interacting with forewing length (FW), as habitat association may depend on the size of the species | $Cr \sim \text{habitat\_sp} * FW + (1 \vert \text{gr}(\text{species\_genetic}, \text{cov} = \text{GenDist})) + (1 \vert \text{species})$ |

## Results

We studied a total of 286 butterfly specimens, of which in total 40 individuals were excluded from the palatability results because 23 trained chicks did not approach either pellet, and 17 chicks consumed less than 50% of the experimental control pellet. The remaining 246 butterfly specimens and 10 mealworm pellets were counted as valid palatability measures, encompassing at least five experimental tests per Eudaminae species (Table 2). Due to the high percentage of non-valid experiments with *Epargyreus spp.* (36%) compared to other species, it is possible that some kind of unpleasant odour is present in this species, although no odour was noted while preparing the pellet.

**Table 2:** Overview of the palatability experiments indicating the number of valid (N) and non-valid experiments per species, average ± SD butterfly body weight, average ± SD morphometric measurements (forewing length, forewing area, wing loading). All values are presented only for valid experiments. Additional morphometric values (forewing width, aspect ratio) and species level sex ratios can be found in Supplement 5.

| Species | N (+ non-valid) | Average body weight [mg] | Forewing length [mm] | Forewing area [mm<sup>2</sup>] | Wing loading [mg/mm<sup>2</sup>] |
| --- | --- | --- | --- | --- | --- |
| *Tenebrio molitor* (Linnaeus, 1758)<br>Palatable reference | 10 (0) | 155.7 ± 76.2* | | | |
| *Heliconius melpomene* (Linnaeus, 1758)<br>Unpalatable reference | 13 (0) | 120.5 ± 25.6 | | | |
| *Cecropterus dorantes* (Stoll, 1790) | 22 (1) | 119.2 ± 24.2 | 22.38 ± 0.87 | 336.05 ± 29.94 | 0.354 ± 0.066 |
| *Cecropterus doryssus* (Swainson, 1831) | 8 (0) | 151.3 ± 29.7 | 21.78 ± 0.80 | 340.75 ± 24.56 | 0.440 ± 0.070 |
| *Cecropterus longipennis* Plötz, 1882 | 31 (8) | 120 ± 26.1 | 19.03 ± 0.56 | 289.16 ± 17.78 | 0.414 ± 0.082 |
| *Cecropterus zarex* (Hübner, 1818) | 5 (1) | 104.2 ± 8.8 | 18.22 ± 0.67 | 260.80 ± 20.87 | 0.402 ± 0.054 |
| *Chioides catillus* (Cramer, 1779) | 17 (2) | 178.6 ± 35.4 | 26.05 ± 1.13 | 425.00 ± 35.19 | 0.418 ± 0.063 |
| *Cogia crameri* (McHenry, 1960) | 13 (1) | 128.5 ± 27 | 22.83 ± 1.03 | 337.08 ± 25.26 | 0.379 ± 0.064 |
| *Drephalys helixus* (Hewitson, 1877) | 7 (0) | 195.7 ± 9.4 | 22.70 ± 0.61 | 322.71 ± 7.57 | 0.607 ± 0.033 |
| *Epargyreus spp.* Hübner, [1819] | 33 (12) | 243.6 ± 32.4 | 28.65 ± 0.92 | 464.54 ± 27.32 | 0.524 ± 0.053 |
| *Spicauda simplicius* (Stoll, 1790) | 27 (4) | 126.9 ± 32.1 | 21.47 ± 0.96 | 332.07 ± 25.27 | 0.381 ± 0.088 |
| *Telegonus fulgerator* (Walch, 1775) | 20 (4) | 343.7 ± 54 | 28.35 ± 1.44 | 507.55 ± 58.13 | 0.678 ± 0.078 |
| *Urbanus esmeraldus / esta* (Butler, 1877) / Evans, 1952 | 11 (1) | 133.1 ± 41.1 | 22.45 ± 1.46 | 337.54 ± 38.69 | 0.395 ± 0.104 |
| *Urbanus spp.* Hübner, [1807] | 11 (1) | 153 ± 43 | 22.56 ± 1.06 | 335.36 ± 28.24 | 0.451 ± 0.103 |

\*Average weight of mealworms per pellet.

Molecular species identification revealed that the sampled *Urbanus* may represent 7 closely related species with low genetic divergences among them, highly similar external morphologies, and similar habitat associations. Due to the low abundances per species (*Urbanus esmeraldus*, n=10; *Urbanus proteus*, n=4; *Urbanus segnestami*, n=2; *Urbanus alva*, n=1; *Urbanus esta*, n=1; *Urbanus belli*, n=1; *Urbanus pronta*, n=1; *Urbanus spp.*, n=2), we considered two strongly supported monophyletic groups as units of analysis in all statistical analyses: one consisting of *Urbanus esmeraldus* and *Urbanus esta* (n = 11) and the other containing the remaining sampled *Urbanus* (n = 11). The COI barcodes of sampled *Epargyreus* have been flagged as potentially representing two species by BOLD (*Epargyreus Burns02* and *Epargyreus Burns06*) but conclusive evidence was not possible to obtain based solely on wing morphology, colour pattern, and habitat use. Thus, *Epargyreus* was analysed as one group. All remaining species have solid identifications based on morphology and COI barcodes (Figure 1).

**Figure 1:**
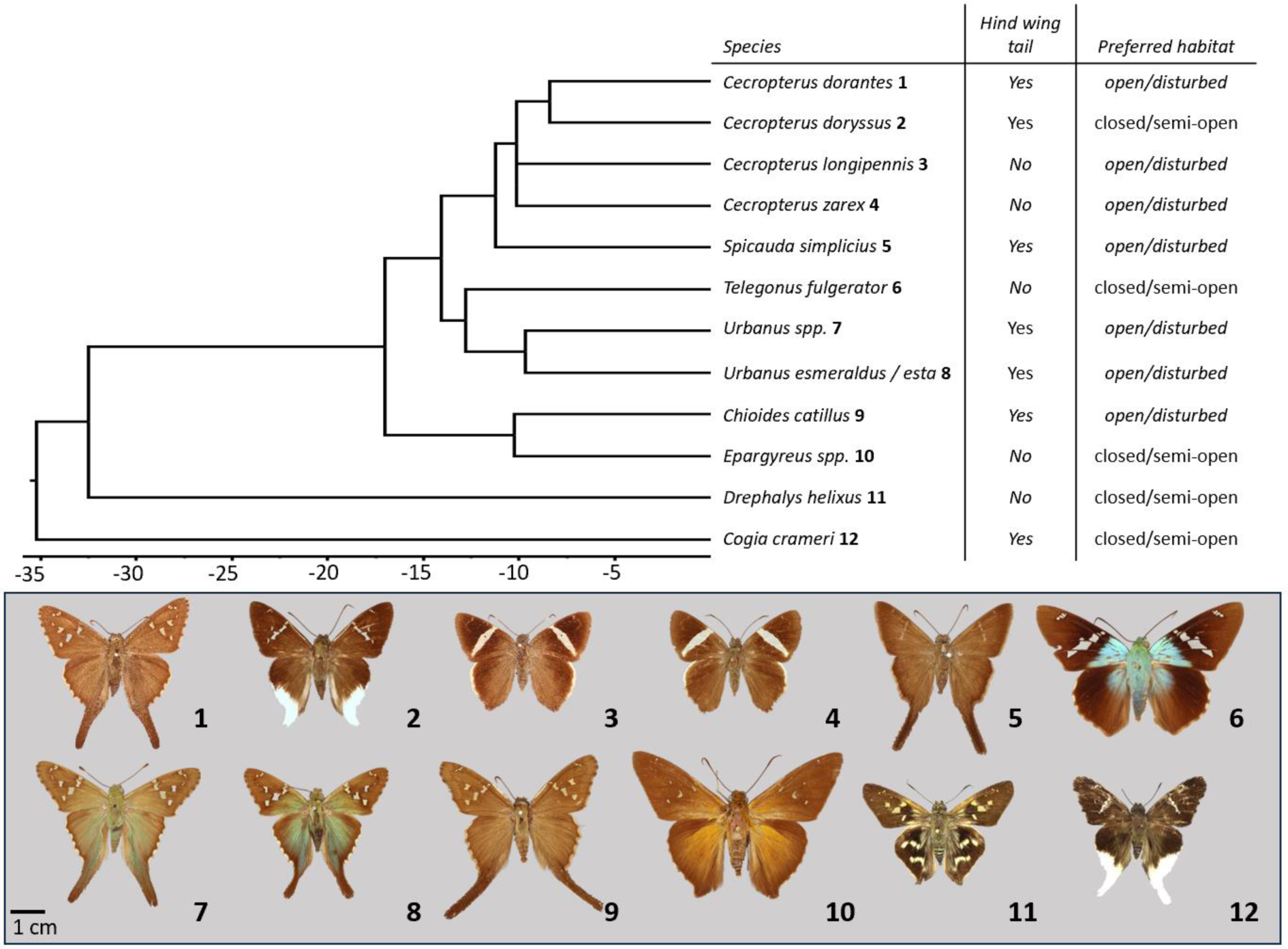
Time-calibrated phylogenetic tree of all sampled Eudaminae species indicating the presence/absence of hind wing tails and the preferred habitats of all species. Butterfly photographs correspond to the species in the phylogenetic tree except, **7** (Urbanus proteus), **8** (Urbanus esmeraldus) and **10** (Epargyreus aspina). Photos taken from specimens collected in Peru and deposited at the Natural History Museum, London, Naturkunde Museum, Berlin, and our collection at the Biology Centre CAS, České Budějovice.

The distribution of consumption ratios (*Cr*, Figure 2) was significantly different from a normal distribution (Shapiro-Wilk normality test, p < 0.0001). The pairwise Wilcoxon signed-rank tests revealed four species being significantly less palatable than our palatable reference (*Tenebrio molitor*), these are *Heliconius melpomene* (the unpalatable reference, p = 0.006), *Telegonus fulgerator* (p =0.004), *Epargyreus spp.* (p = 0.002) and C*ecropterus longipennis* (p = 0.026). Additionally, three species were almost significantly less palatable than *Tenebrio molitor*, these are *Chioides catillus, Drephalys helixus and Urbanus spp.* (p = 0.077, p = 0.069 and p= 0.075, respectively). *Cecropterus longipennis* was the only Eudaminae species close to being significantly different from *Heliconius melpomene* (unpalatable reference, p = 0.075). While the unpalatable and palatable references showed low intra-group variability, we detected a remarkable high amount of inter- and intra-specific variability across all tested Eudaminae species. Further, we were not able to reject the null hypothesis that palatability does not exhibit phylogenetic signal, as using species level *Cr* averages and medians yielded mixed results for both, Pageĺs lambda (species-level average *Cr*: λ = 1.25, p = 0.29; median *Cr*: λ < 0.001, p = 1) and Bloomberg’s K (average *Cr*: K = 0.93, p = 0.045; median *Cr*: K = 0.77, p = 0.13). However, our limited phylogenetic sampling coupled with the high variability within *Cr* hinders any conclusive interpretation regarding the evolution of palatability within Eudaminae.

**Figure 2:**
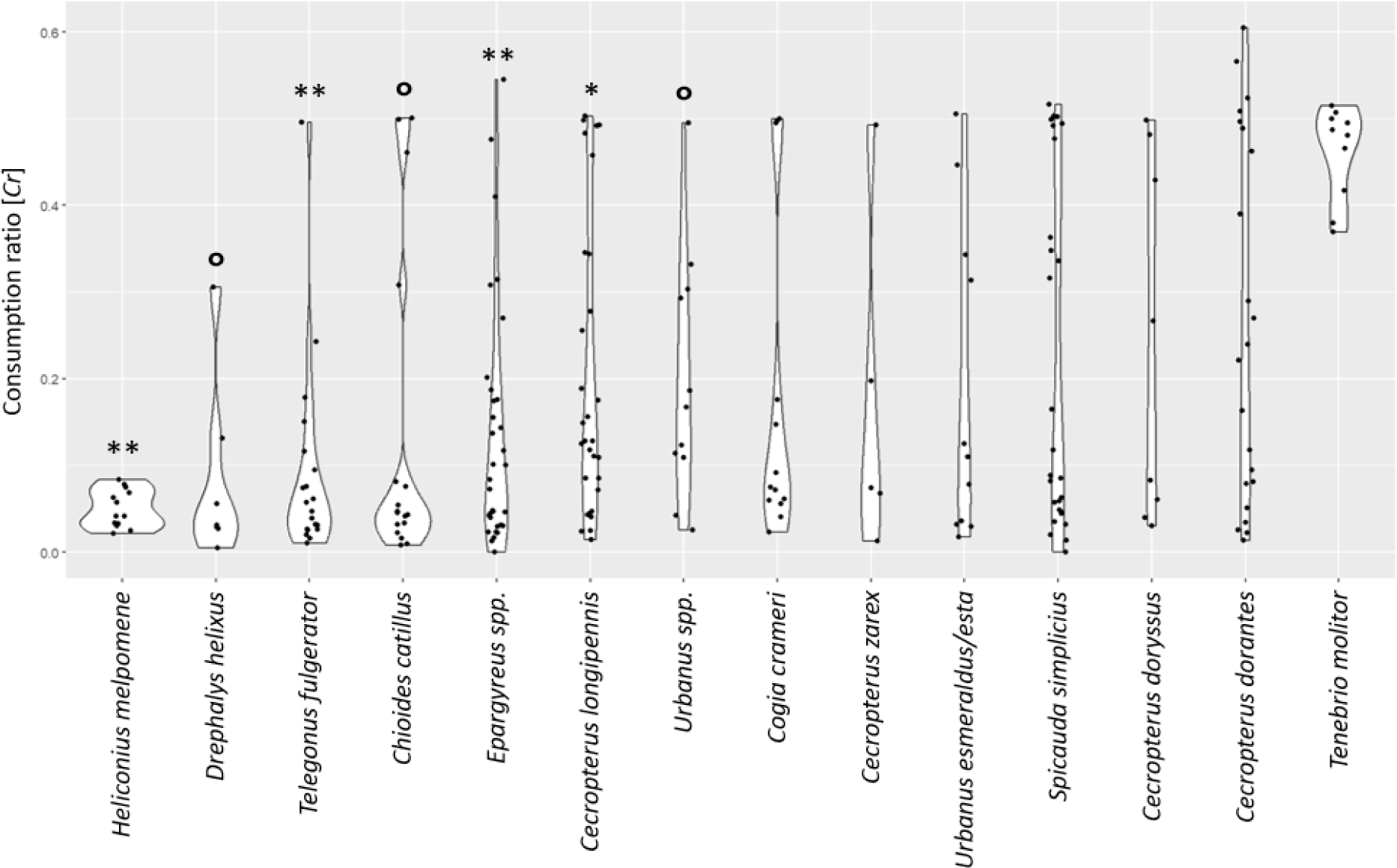
Consumption ratios (Cr) of all abundant Eudaminae species at the study locality. Heliconius melpomene was used as the unpalatable reference and Tenebrius molitor as the palatable reference. Asterisks (*) positioned above the plots denote significant differences in Consumption Ratio (Cr) compared to the palatable references, as determined by pairwise Wilcoxon signed-rank tests (* = 0.01<p<0.05, ** = 0.001<p<0.01), while ° signifies a p-value approaching significance (below 0.08).

The Bayesian MCMC regression analyses resulted in Rhat values (potential scale reduction factor, Gelman-Rubin statistic) consistently at 1 and effective sample sizes (ESS) not lower than 3000. The full output from all presented models can be found in Supplement 6. A plot for the difference in palatability between male and female skippers can be found in Supplement 7.

When testing for correlations between hindwing tails and wing loading (prediction**(I)**, Figure 3-A), we found a significant negative correlation (mean = −0.25, SD = 0.12, L-95% = −0.50, U-95% = −0.01), which goes against our initial prediction. This result indicates that butterflies with high wing loading (estimate of increased flight speed) are usually lacking hind wing tails.

**Figure 3:**
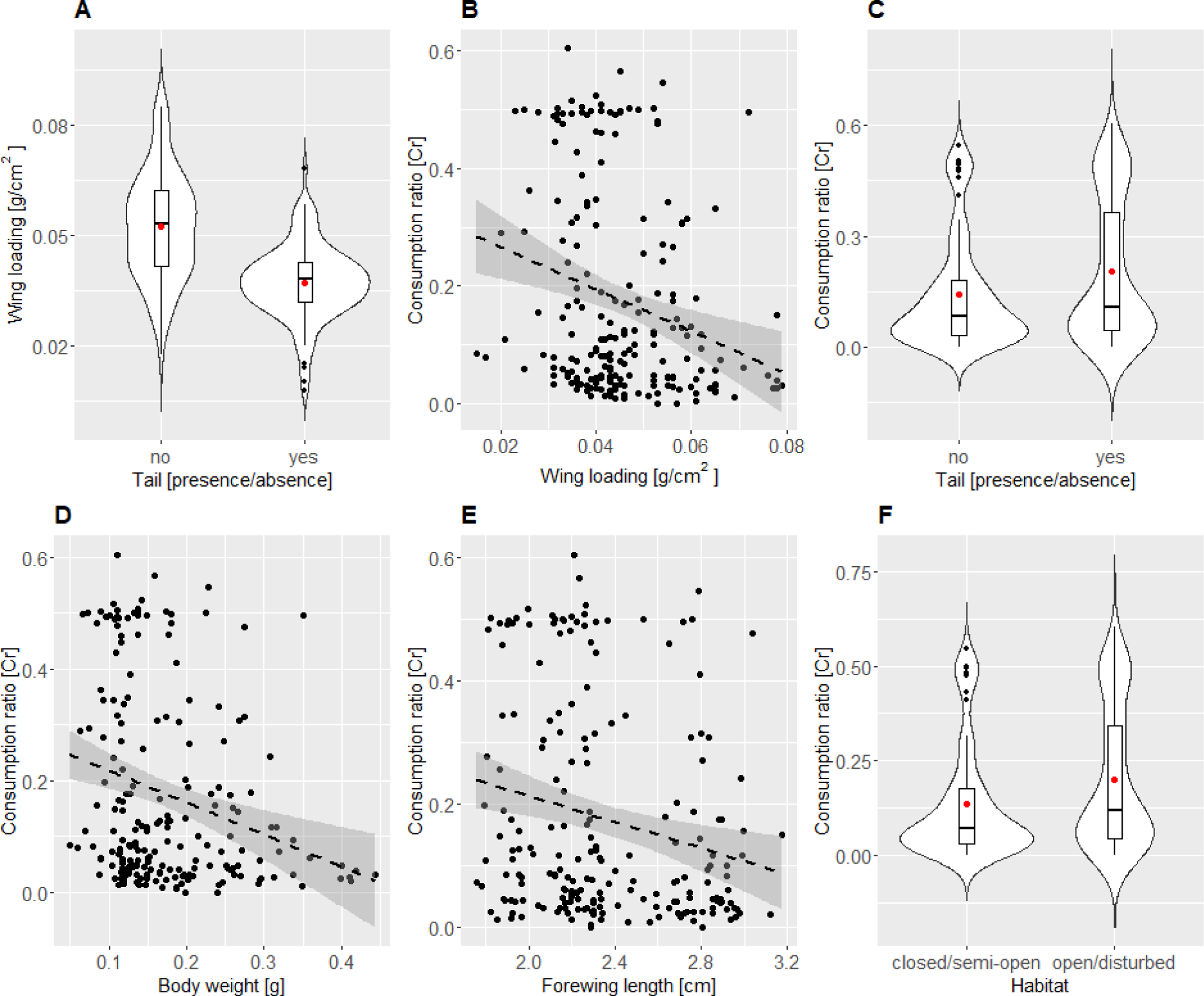
Depictions of all models used to test initial predictions: **A** Comparison of species with or without hindwing tails in relation to wing loading (positively correlated with flight speed); **B** Linear regression between consumption ratio (Cr, measure for palatability) and wing loading; **C** Violin plot comparing Cr between species expressing hindwing tails and species without hindwing tails; **D** Linear regression between total body weight and Cr; **E** Linear regression between forewing length and Cr; **F** violin plot comparing wing loading between species found predominantly in closed/semi-open habitats or open/disturbed habitats. None of the graphs includes data from the palatable or unpalatable references. Red dot in violin plots represents the mean.

Contrary to our initial predictions, we found a significant positive effect of wing loading on *Cr* values (prediction **(II)**, Figure 3-B, mean = −19.53, SD = 6.27, L-95% = −31.93, U-95% = −6.91) (Figure 3-A), indicating that specimens with high wing loading (e.g., *Telegonus fulgerator, Epargyreus spp.*) are less palatable than butterflies with low wing loading (e.g., *Cecropterus dorantes*, *Spicauda simplicius*).

When estimating the effect of hindwing tails (mechanical defence) on palatability (prediction **(III)**, Figure 3-C) while accounting for its interaction with forewing length (the functional effect of tails depending on the forewing size), we did not detect any significant correlation (mean = 1.71, SD = 1.57, L-95% = −1.35, U-95% = 4.87). Which implies, that contrary to our initial prediction hindwing tails occur in both unpalatable and palatable Eudaminae butterflies.

When testing the influence of body size (body weight and forewing length, separately) on *Cr* (prediction **(IV)**, Figure 3-D/E), we also recovered significant negative effects, following our initial prediction, although with lower effect than that of wing loading (mean = −3.16, SD = 1.08, L-95% = −5.30, U-95% = −1.06, for body weight; mean = −0.57, SD = 0.26, L-95% = −1.09, U-95% = −0.05, for forewing length).

Regarding prediction **(V)** (Figure 3-F), we found no correlation between habitat association and *Cr* values, thus, consumption deterrence driven by reduced palatability (potentially a chemical defence) in Eudaminae is not associated with species predominantly occurring in either closed or open environments as initially predicted. The influence of forewing length, as well as its interactions with habitat was not significant.

The effects of phylogenetic and non-phylogenetic species-level variance in all models were insignificant and low (∼0.03 and ∼0.12, respectively).

## Discussion

By quantifying individual- and species-level traits related to anti-predator defences, we revealed multi-defence strategies used by abundant skipper butterflies co-occurring at the same locality. These defences encompass behavioural aspects (where high wing loading predicts increased flight speed), mechanical features (such as hindwing tails), and potentially chemical elements (resulting in less palatability for surrogate predators), which often appear in different combinations among species. Strikingly, we unveiled a broad range of palatability levels between and within skipper species, from almost as unpalatable and with low intra-specific variation as *Heliconius melpomene* (e.g., *Drephalys helixus*, *Telegonus fulgerator* and *Chioides catillus*) to almost as palatable as mealworms but apparently with high intra-specific variation (e.g., *Cecropterus dorantes* and *S. simplicius*). Our results bring substantial evidence that seemingly unpalatable butterflies can simultaneously rely on other rarely reported behavioural (fast flight) and mechanical defences (brittle structures that deflect attack as hindwing tails). These findings counterbalance the commonly reported combinations of defences in prey, where chemical defence is often associated with slow movement or morphological resilience (Kikuchi et al., 2023; but see Pinheiro, 1996). Additionally, unpalatability seems linked to increasing body size (both regarding body weight and forewing length).

It is not clear whether consumption ratio (*Cr*), a measure of palatability, harbours phylogenetic signal, but future studies with increased phylogenetic sampling will provide more robust insights into the evolution of (un)palatability in Eudaminae. The limited host plant records for Eudaminae (Janzen & Hallwachs, 2009; Robinson et al., 2023) poses challenges to understand the potential sources of their unpalatability. Faboideae legumes (family Fabaceae) are typical host plants, with quinolizidine (Misra et al., 1999) and pyrrolizidine alkaloids (Cogni & Trigo, 2016; Irmer et al., 2015) reported in a few species, though these metabolites are generally rare across Fabaceae. Pyrrolizidine alkaloids are exploited as a host plant-derived chemical defence by unpalatable Ithominii and Danainae butterflies (Trigo et al., 1996). Eudaminae butterflies that are less palatable might potentially employ metabolites from Faboideae, although this remains speculative. Definitive conclusions await further documentation of host plants or additional experiments investigating the presence and quantity of secondary compounds in adult skippers.

Adult Eudaminae skippers do not display typical visual defences such as warning colourations and aposematism associated with unpalatable Lepidoptera (Kikuchi et al., 2023). However, if host plant-derived chemical defence is confirmed in other Eudaminae species, we cannot rule out that unpalatability is associated with the highly aposematic caterpillars of some Eudaminae (e.g., *Telegonus fulgerator*: Hebert et al., 2004; Janzen & Hallwachs, 2009 or *Chioides catillus*: Cock, 2016), which might indeed represent honest unprofitability signals and might be widespread in Eudaminae. This suggests the possibility that unpalatability in Eudaminae skippers may have been primarily selected during the larval, less mobile, stage, as found in other Lepidoptera species (Poloni et al., 2023), while still providing some protection during the adult stage. Therefore, while adult Eudaminae likely rely on fast flight and escape ability, a feature that may have been inherited from their common ancestor (the family Hesperiidae, in general, have highly erratic and fast flights), unpalatability may not be critically under selection in adults but may also not be strongly selected against, explaining partially its high intra- and interspecific variability.

The high intra-specific variability in palatability levels within Eudaminae might be the result of relaxed selective pressures, in contrast to highly unpalatable butterflies like *Heliconius*. Skippers are difficult to capture (behavioural defence) and possess deflective structures on their wings (mechanical defence), thus covering anti-predator defences acting at different stages of the predation sequence. If Eudaminae phenotypes effectively signal their evasive ability to potential predators (visual defence; e.g., Linke et al., 2022), then unpalatability might not play a major role as anti-predatory defence in adult Eudaminae as in other highly aposematic and unpalatable butterflies, but a complementary role together with other types of defences. Therefore, adult skippers, with their large thoracic muscles, might be still of high nutritional value and profitable to chase for specialized insectivorous predators, but very likely highly unprofitable to other flying predators.

Additionally, we found significant correlational evidence that unpalatability is explained by increasing wing loading. Due to time and logistics limitations in the field, we could not directly measure flight behaviour and performance of the studied species. Thus, we rely on a substantial body of evidence suggesting that wing loading is positively associated with high flight performance and speed in butterflies (Dudley, 1990; Srygley & Chai, 1990; Dudley & Srygley, 1994; Berwaerts et al., 2002; Le Roy et al., 2019), to propose that certain Eudaminae species likely combine behavioural (fast flight) and consumption deterrence defences (decreased palatability). One plausible explanation is that as wing loading increases, there is a simultaneous increase in body mass and/or a decrease in wing area.

Consequently, these species may be more energetically profitable for specialized predators, prompting such prey to benefit from the expression of multiple anti-predator defences (Pinheiro et al., 2016). Our observation that multi-defences in prey are size dependent is unlikely to result from our experimental design. Even when presenting large pellet sizes during the experiments, starving chicks were eager to consume more than two pellets and to keep actively foraging in the experimental box (chick pre-training observations). Therefore, larger butterflies seem to be more chemically defended and might further increase their survival probability by also having highly evasive flights (e.g., as seen in some species of *Battus* and Heliconiini butterflies, Pinheiro et al., 2016), which is a defence combination rarely reported for Lepidoptera (Kikuchi et al., 2023).

Mechanical defences such as hindwing tails divert attacks towards non-vital body parts, and are easily breakable (Chotard et al., 2022). They are usually co-expressed with behavioural defences such as fast flight, as is the case in Eudaminae. However, breakable parts are commonly absent in unpalatable butterflies such as *Heliconius* and Ithomiini, as well as other Lepidoptera (Kikuchi et al., 2023). We did not find evidence for such a negative correlation in Eudaminae because hindwing tails can be present in both, largely palatable or largely unpalatable species. The most noteworthy example is the generally unpalatable *Chioides catillus* which possesses long hindwing tails. In this case, we propose that such a dual defence is also driven by their large body size (body weight and forewing area), which might leave them prone to higher predation due to increased detectability and/or predator preference. Interestingly, hindwing tails appear more frequently in species with intermediate forewing length, potentially indicating a functional trade-off between size and attack deflection (both for larger and smaller individuals). In smaller species, the distance between hindwing tails and the body may be too small to effectively deflect predator attacks, rendering hindwing tails insignificant, as observed in species like *Cecropterus longipennis* and *C. zarex*. Finally, we cannot rule out that hindwing tails might as well serve as visual defences, thus, representing a multi-role trait that acts early (visual defence – identification) and late in the predation sequencing (mechanical defence – approach) (Broom et al., 2010; Kikuchi et al., 2023). In this regard, Linke et al. (2022) reported that hindwing tails might be learnt as cues signalling unprofitability, such as evasive ability, or in, *Chioides catillus*, unpalatability.

In addition to the hindwing tails, we gathered evidence that Eudaminae wing structures or chemical compounds in the wings may influence their palatability (Supplement 4). Skippers have abundant hair-like scales on their wings, which may be mechanically unpleasant to handle by certain avian predators. Alternatively, chemical compounds present within wing tissue, as in some Ithomiini butterflies (McClure et al., 2019), may be behind the apparent increased unpalatability of *Spicauda* wings. Furthermore, in general females appear to be less palatable than males (see Supplement 7), but we remain cautious about this result as the sample size was low. However, if the pattern holds with increased sampling size, females may rely on stronger unpalatability defence as they and their eggs would benefit from being released alive by naïve predators upon smell and taste. Alternatively, due to their enlarged abdomens to carry the eggs, their evasive ability may be compromised, and becoming more unpalatable may compensate their anti-predator strategies, similar to female-limited Batesian mimicry as in *Papilio polytes* (Nishikawa et al., 2015).

We did not find any significant correlation between unpalatability and habitat occurrence, as predicted by Srygley & Chai (1990). Although our dataset seems to have small explanatory power, it is worth noticing that species found on hill tops tended to be relatively large (pers. observation D. Linke, P. Matos-Maraví), fast (high wing loading), and unpalatable (e.g., *Epargyreus* spp. and *Drephalys helixus*). While large butterflies are typically found on tropical hill tops (Henriques et al., 2022), the predicted fast flights of Eudaminae on the hilltop environment might be explained by the required powered flight due to more windy conditions compared to other habitats in the study location. Alternatively, the bird community in this habitat might differ from nearby forest and open environments, potentially exposing them to high predation pressure from agile insectivorous birds. Thus, while potentially being more rewarding for predators due to their high body mass, their high unpalatability may further increase their chances of survival. Interestingly, despite their deep evolutionary divergence, the phenotypes of *Epargyreus* and *Drephalys* seem to resemble also those in other Neotropical localities (Burns & Janzen, 2000). Thus, mimicry driven by either unpalatability or escape ability (Janzen et al., 2009; Páez et al., 2021; Linke et al., 2022) is possible in these groups, though further evidence is needed from other *Epargyreus* and *Drephalys* communities.

Our findings uncovered butterfly species that use multiple defence strategies rarely reported in the literature (e.g., reduced unpalatability, fast flight, and brittle structures such as hindwing tails), but to characterize their evolution, including directionality and transition rates, future efforts are needed to unveil larval diets and chemical profiles, as well as a thorough quantification of flight behaviours across lineages. Nevertheless, our study adds further evidence that coupling multiple anti-predator strategies might not be uncommon among butterflies and moths (Pinheiro & Freitas, 2014; Pinheiro et al., 2016). Indeed, having multiple anti-predator defences may be better than having just one (Kang et al., 2017), and in Eudaminae, it seems to be determined by prey detectability and/or preference (explained by body size) rather than habitat associations, though the latter seems to be explained by wing loading (fast flight).

## Acknowledgments

We would like to thank all persons helping us during the experiments in the field, especially César Ramírez and Stephanie Gallusser who allowed us to conduct the experiments in their properties and were exceptionally helpful during field work. We would also like to thank Eva Kriegová and Barbora Bělovská for their help in the molecular laboratory. Additionally, we would like to extent our gratitude towards Leonardo Ré Jorge for his help and advice in the statistical analysis.

## Data availability

The data that support the findings of this study are available from the corresponding author upon reasonable request and will be archived on Zenodo after acceptance. Barcodes generated by this project will be deposited in BOLD and / or GenBank.

## Authors contributions

**DL** – study planning, field investigation, data curation and evaluation, writing original draft, review and editing; **JHM** – field investigation, review and editing; **VNPNE** – field investigation, review and editing; **PGR** – laboratory work, review and editing; **LSS** – review and editing; **ME** – study planning, review and editing; **PMM** – study planning, field investigation, funding acquisition, review and editing.

## Funding

Funding was provided by the Junior GAČR grant (GJ20-18566Y) and the PPLZ program of the Czech Academy of Sciences (fellowship grant L20096195) as well as GAJU n.014/2022/P. Photography in the Natural History Museum (NHM) in London was funded by SYNTHESYS+ under GB-TAF-TA4-015. Logistic support in Peru was supported by 421 Fundación San Marcos, Letty Salinas was partially funded by Universidad Nacional Mayor de San Marcos RR 05557-R-22 (Project B22100321).

Computational resources were provided by the e-INFRA CZ project (ID:90140), supported by the Ministry of Education, Youth and Sports of the Czech Republic.

Butterflies were collected under the national permits RDG N.° 0488-2019-MINAGRI-SERFOR-DGGSPFFS, D000088-2021-MIDAGRI-SERFOR-DGGSPFFS, and D000047-2022-MIDAGRI-SERFOR-DGGSPFFS-DGSPFS and exported under the permits 003735-SERFOR and 003758-SERFOR.

